# Stimulated *Prorocentrum donghaiense* cell growth by *in-situ* mariculture dissolved organic matter

**DOI:** 10.1101/2024.02.02.578562

**Authors:** Hongwei Wang, Siyang Wu, Jian Ma, Yiting Hong, Chentao Guo, Jing Zhao, Xin Lin

## Abstract

Mixotrophic dinoflagellates frequently cause harmful algal blooms (HAB) in eutrophic mariculture waters that contain diverse excreted dissolved organic matter (DOM). The phagotrophy and the utilization of single organic small molecules have been extensively investigated in the bloom-forming mixotrophic dinoflagellates. However, their ability to utilize the *in-situ* DOM via absorbtrophy still remains unexplored. Here we examined the growth promotion effect of the *in-situ* mariculture DOM on *Prorocentrum donghaiense*, a representative HAB-forming species in coastal waters. Our results showed that the cell growth and photosynthesis of *P. donghaiense* were significantly promoted under *in-situ* DOM culture conditions. Additionally, parallel cultures were set up to disclose the potential role of the bacterioplankton in the free-living community (helper), where they aid in the remineralization of the *in-situ* DOM, and the phycosphere community (competitor), where they compete against the algal host to acquire nutrients from the *in-situ* DOM. Meanwhile, we determined the cellular stoichiometry under different culture conditions, showing that mariculture DOM can shape cellular stoichiometry significantly. Elevated cellular N (84.96%) and P (48.3%) were observed in spring DOM groups compared with the control groups. For the first time, this study quantifies the efficient utilization of the *in-situ* DOM via absorbtrophy, indicating the vital role in the outbreak and maintenance of HAB events.

## 1. Introduction

Harmful algal blooms (HABs) are characterized by rapid proliferation and accumulation of algae and protists in aquatic ecosystems, and have been a global concern for decades due to the negative effects on coastal ecosystems and public health (GlobalHAB, 2017). Eutrophication, caused by excess anthropogenic nutrient inputs, is responsible for the frequent outbursts of HAB events in coastal waters (Glibert, 2017; Malone and Newton, 2020). Rapid expansion of mariculture is believed to be one of the major contributors to eutrophic coastal waters, as on the southeast coast of China (Wang et al., 2020a; Li et al. 2021a).

Mariculture generates extra nutrient inputs through a variety of ways, including bait residue, metabolic wastes and feces from cultured species, and remineralization of microorganisms. Most baits have a high protein content, but the nutrient retention efficiency of reared organisms is inadequate, averaging less than 35% (Bouwman et al., 2013; Cao et al., 2015; Meng and Feagin, 2019). As a consequence, for example, nitrogen concentration can reach as high as 39.6-87.5 µM in the Sansha Bay mariculture area in winter (Han et al., 2020). A considerable portion of the waste is released in dissolved form, while particulate matter is also decomposed into various dissolved compounds via biological metabolism, leading to a complex composition of DOM in the aquaculture environment (Bouwman et al., 2013; Yang et al., 2017) and affecting the cycling of key elements (Xiao et al., 2023). Advanced chemical analysis reveals that DOM is found to be highly enriched in mariculture waters, with DOC, CDOM and FDOM significantly higher than the adjacent waters (Yang et al.,2022). Under such circumstances, DOM may serve as extra nutrient sources to support rapid cell growth and lead to the outbreak of HABs (Bouwman et al., 2013).

Mixotrophic dinoflagellates are major groups of HAB-causative organisms, accounting for 40% of the species forming HABs globally (1990-2019) (Jeong et al., 2021). In the East China Sea (ECS), mixotrophic dinoflagellates have dominated bloom events in recent years, and *P. donghaiense* is the typical HAB causative species in ECS mariculture area (Spilling et al. 2018; Li et al., 2021b). The mixotrophic strategies adopted by *P. donghaiense*, including the phagotrophy and utilization of single small molecular compounds, have been well investigated in lab (Lin et al., 2016; Shi et al., 2017). The mixotrophic ability confers important competitive strategy for dinoflagellates, and has been reckoned critical for them to prevail in the phytoplankton community and form HABs (Stoecker et al., 2017; Jeong et al., 2021).

As a major mechanism of mixotrophy, absorbotrophic organisms uptake soluble organic matter through active uptake or/and pinocytosis to obtain carbon and nutrients (Flynn et al., 2013, Schmidt et al., 2013). Increasing evidence shows that algae can utilize a wide range of carbon substrates (e.g., acetate, ethanol, and fatty acids, etc.) (Selosse et al., 2017). A couple of dinoflagellates can directly uptake macromolecules via pinocytosis (Granéli et al., 1999), and absorbotrophic uptake of DMSP by the heterotrophic dinoflagellate *Oxyrrhis marina* has also been demonstrated experimentally (Saló et al., 2009). Absorbotrophic mixotrophy is commonly found in organic matter-rich meso-eutrophic waters and have been implicated as a driver of massive harmful algal blooms (Burkholder et al., 2008; Selosse et al., 2017).

However, the utilization of *in-situ* DOM by mixotrophic dinoflagellate still remain unexplored. To address this issue, we use *in-situ* abalone (a large-scale, typical breeding species in the algal bloom-prone areas of Fujian, China) mariculture seawater as the sole medium to grow *P. donghaiense*. We firstly examined the effect on cell physiological responses between filtered seawater (DOM contained) sampled in spring and autumn respectively. Meanwhile, pairwise experimental groups were set up to compare the function of bacterioplankton between *in-situ* and phycosphere communities regarding the DOM utilization. Cellular stoichiometry is determined to explore the contribution of biogeochemical elements in DOM to algal growth, and the ecological implications are discussed.

## 2. Materials and Methods

### 2.1 DOM filtration, culture condition and experiment set up

*In-situ* seawater was collected in April (spring) and October (autumn) from the abalone mariculture area in Dongshan County, Fujian Province. In this study, the DOM culture refers to the culture medium prepared with the filtrate of the collected seawater, which was filtered through a 0.22 µm pore size membrane three times (Carlson and Hansell, 2015). Batch cultures were set up to investigate the effect of DOM with different treatments, which is summarized in Fig. 1. Filtrate (∼20L) acquired from immediate filtration of *in-situ* seawater was appointed as the initial DOM (iDOM). Meanwhile, another volume of *in-situ* seawater (∼20L) was kept in the dark at room temperature for 20 days for further microbial degradation of DOM, and the filtrate afterwards was appointed as mDOM.

**Fig. 1.**
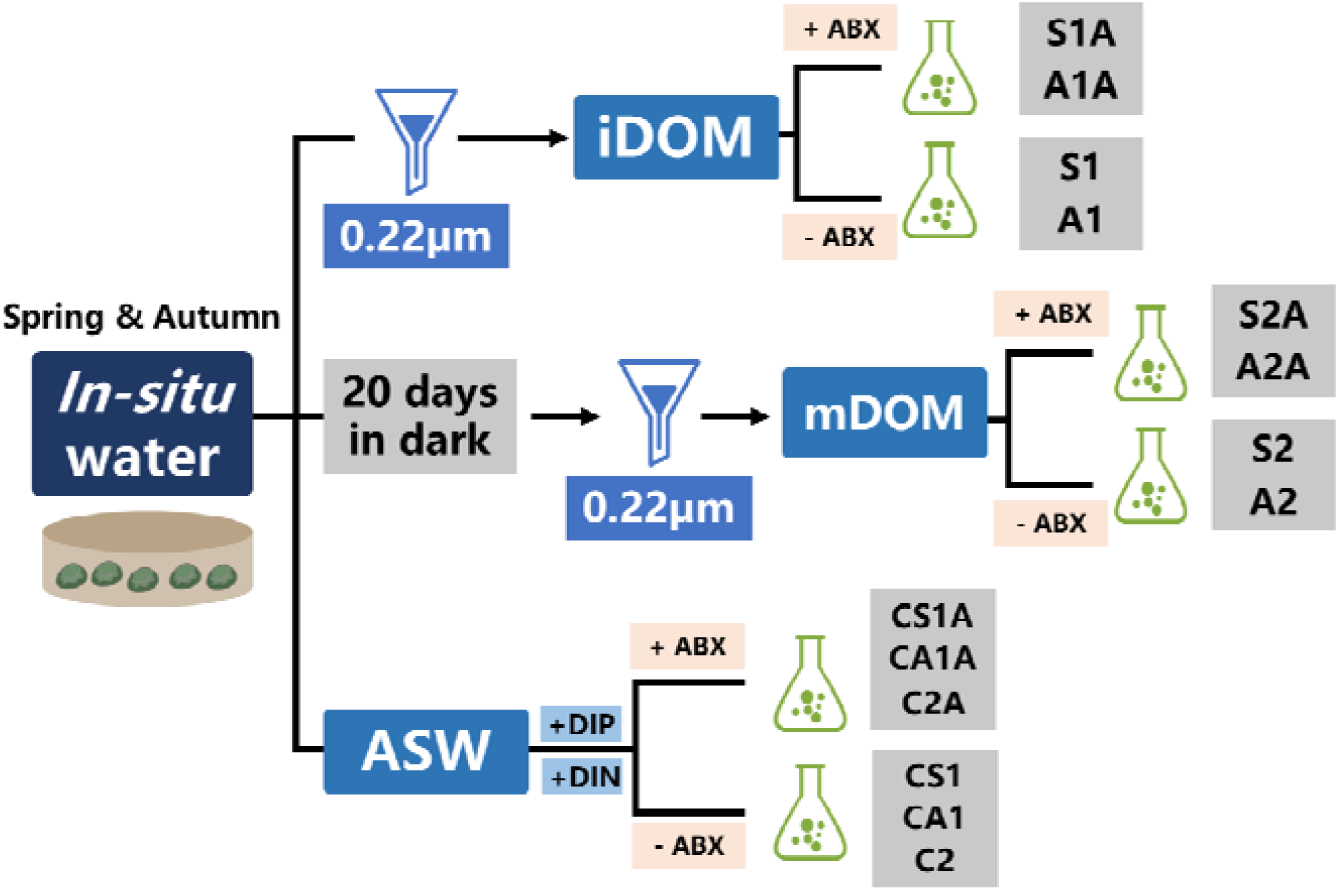
*In-situ* DOM filtration and experiment design. iDOM: initial DOM; mDOM: microbial degraded DOM; ABX: antibiotics; ASW: artificial seawater.

*P. donghaiense* was kindly provided by the Collection Center of Marine Algae, Xiamen University, China. Seed culture was grown in the oceanic waters (average concentration of DIN and DIP less than 1µM) for 2 weeks to deplete intracellular nutrient storage. To mimic the *in-situ* scenario, filtrate was used as the sole medium substrate, and the initial cell concentration was 80 cells mL^-1^ referred to reported pre-bloom concentration (Ministry of Natural Resources of the People’s Republic of China, 2021). The control group was grown in the medium prepared in artificial seawater (Sunda et al., 2005), in which DIN and DIP were provided according to the measured values of the filtrate respectively (Supplementary Table 1). To examine the effect of phycosphere bacteria, we set up two parallel groups for each condition. One group was treated with the antibiotic cocktail KAS (final concentration in the culture, 100 µg/mL ampicillin, 50 µg/mL streptomycin, and 50 µg/mL kanamycin) and the other was not.

Cultures were incubated at 20 D under a light: dark cycle of 14: 10 h with a photon flux of 100 µE m^-2^ s^-1^. Different groups are labeled as the following, S/A (spring/autumn)-1/2 (i/mDOM)-A (antibiotic) (Fig. 1, Supplementary Table. 1).

### 2.2 Cell concentration, chlorophyll-a (Chla) content and *Fv/Fm*

Cell concentration, Chla and photochemical efficiency *Fv/Fm* were measured every two days during the course of the experiments. Cell concentration was determined by cell count using a Sedgewick-Rafter counting chamber (Wildco, MI, USA). Specific growth rate (*µ*) was calculated as the following equation, *µ* = ln(*N_2_*/*N_1_*) / (*t_2_* - *t_1_*), where N_2_ and N_1_ represent cell concentration at the time points 2 (t2) and 1 (t1), respectively. To determine cellular Chla content, 25 mL of cell culture were filtered onto a 25mm GF/F membrane and then immersed in 94% acetone over 24 h in the dark at 4 D. After centrifugation at 3500 rpm for 2 min, the Chla content in the supernatant was measured using Turner Trilogy (Turner Designs fluorometer, USA). Another cell sample (2 mL) was collected and shielded from light for 20 min, then the photochemical efficiency of photosystem II *Fv/Fm* was quantified by FIRe Fluorometer System (Satlantic, Halifax, NS, Canada) in the dark.

### 2.3 Dissolved inorganic nitrogen (DIN) and dissolved inorganic phosphorus (DIP)

A 15mL sample was removed from each culture and filtered through a GF/F membrane, and the filtrate was used to measure DIN (sum of nitrate, nitrite, and ammonium) and DIP concentration using the autonomous portable analyzer—*i*SEA-II (Xiamen university, China) (Fang et al., 2022).

### 2.4 Particulate phosphorous, cellular carbon and nitrogen contents

For particulate phosphorus (PP) measurement, 25 mL of the cell culture were filtered onto a 25mm GF/F membrane previously combusted at 450D for 5 h. The membrane was then placed in a 30 mL glass digestion tube, adding 25 mL MiliQ water and 4 mL 5% Potassium persulfate solution, and digested in an autoclave at 121 D for 0.5-1h. Digested sample was filtered through GF/F membrane and the filtrate was used to measure PP content using phosphorus molybdenum blue spectrophotometry (Jeffries et al., 1979; Murphy and Riley, 1962). The total PP content was divided by the cell number in the sample to convert to per cell content.

Cellular carbon and nitrogen contents were determined using a vario EL cube analyzer (Elementar Analysensysteme GmbH, Hanau, Germany). 25 mL of the cell culture were filtered onto a 25mm GF/F membrane previously combusted at 450D for 5h and then the membranes were stored at –80°C. Before determinations, the membranes were dried at 56 D for 24 h, then 500 µL of 1% HCl was dripped onto the membrane, followed by drying at 56 D for another 24 h. After that, the filtered membranes were wrapped with tin foil and then compressed into small pellets of uniform size for subsequent elemental analysis. The carbon and nitrogen content were calculated as per cell, and C/N ratios were obtained.

### 2.5 Statistical analysis

All data presented are means with standard deviation calculated from the triplicated cultures in each group. We applied the software SPSS ver. 26.0 to evaluate the statistical significance of the differences observed among groups at the level of *P* < 0.05.

## 3. Results

### 3.1 Promoted cell growth by *in-situ* DOM

The initial concentration of inorganic nutrients varied among experimental groups (Supplementary Table. 1). Thus, the matching concentration of DIN and DIP was provided in the control group (Supplementary Fig. 1f). Starting with a same cell density of 80 cells mL^-1^, *P. donghaiense* cells cultured with iDOM outgrow the control group after 5 days, but it differed in the mDOM groups (Fig. 2). In the later stages of the incubation, cell density in most of the DOM culture groups was significantly higher than that of the corresponding control groups (T-test, *P* < 0.05) (Fig. 2 a, b, c), indicating that DOM can be utilized as an extra source of nutrients to support cell growth, with the exception in the autumn-mDOM groups (Fig. 2d). The average growth rates during exponential phase of the DOM culture groups ranged from 0.326-0.447 day^-1^, much higher than that of control groups of 0.122-0.262 day^-1^ (Fig. 2e) (ANOVA, *P* <0.05). Consistent with cell growth, DIP and DIN in the medium decreased gradually over time and were depleted when culture entered the plateau phase (∼ day 10), except for spring-DIP (Supplementary Fig. 1a-d).

**Fig. 2.**
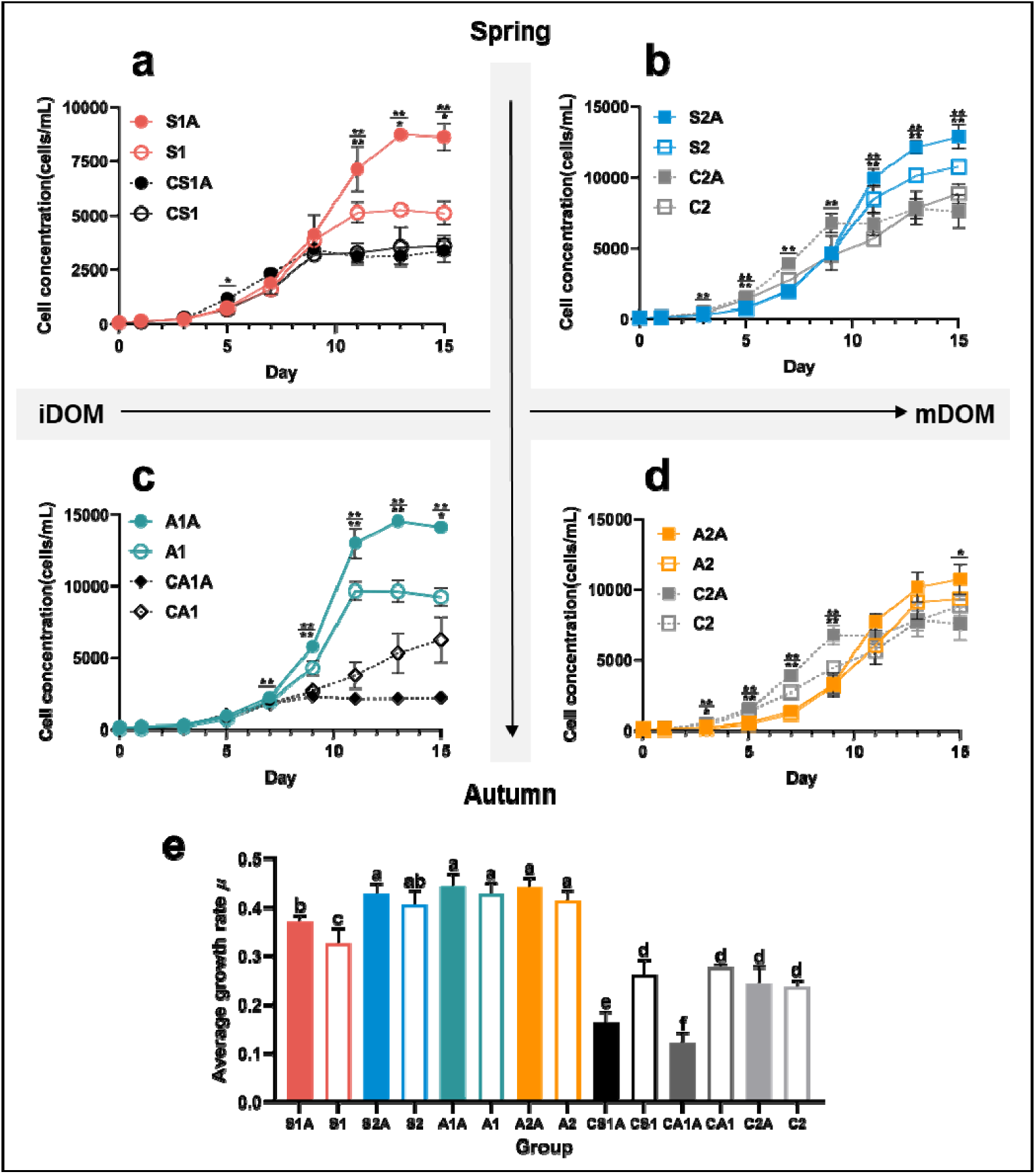
Cell growth under different DOM conditions. (a) Growth curves under spring-iDOM conditions. (b) Growth curves under spring-mDOM conditions. (c) Growth curves under autumn-iDOM conditions. (d) Growth curves under autumn-mDOM conditions. (e) Average growth rate (µ) of each group during the exponential growth phase (Day 5 ∼ Day 11).

Whilst, the addition of antibiotics against phycosphere bacterial community posed positive effect on the cell growth in most cases (Fig. 2). With the antibiotics treatment, the highest value of the cell density was acquired in the A1A group, 14561 cells mL^-1^ of day 13, which was 6.5-fold change of the CA1A, and 2.6-fold change in the S1A group against CS1A comparably. Whereas in non-antibiotic groups, the elevation of cell growth was limited with 1.5- fold change (S1 vs. CS1) and 1.6- fold change (A1 vs. CA1). (Fig. 2a, c) Besides those, the cell density in the CA1A group was 43.2% of that in the CA1 when grown with the supply of the mimic inorganic nutrient (Fig. 2c).

Consistent with cell growth, *Fv/Fm* representing the photosynthetic capacity and cellular Chla content exhibited the similar trend (Fig. 3). In DOM cultures, *Fv/Fm* maintained at 0.656-0.760 after the 9th day, compared to 0.497-0.650 in the control groups (Fig. 3a, b, c, d) (T-test, *P* <0.05). The cellular Chla during exponential phase of DOM groups also showed a notably increase to 0.764 - 1.715 pg/cell, 32.37% - 70.71% higher than the control groups (Fig.3 e, f) (T-test, *P* <0.01).

**Fig. 3.**
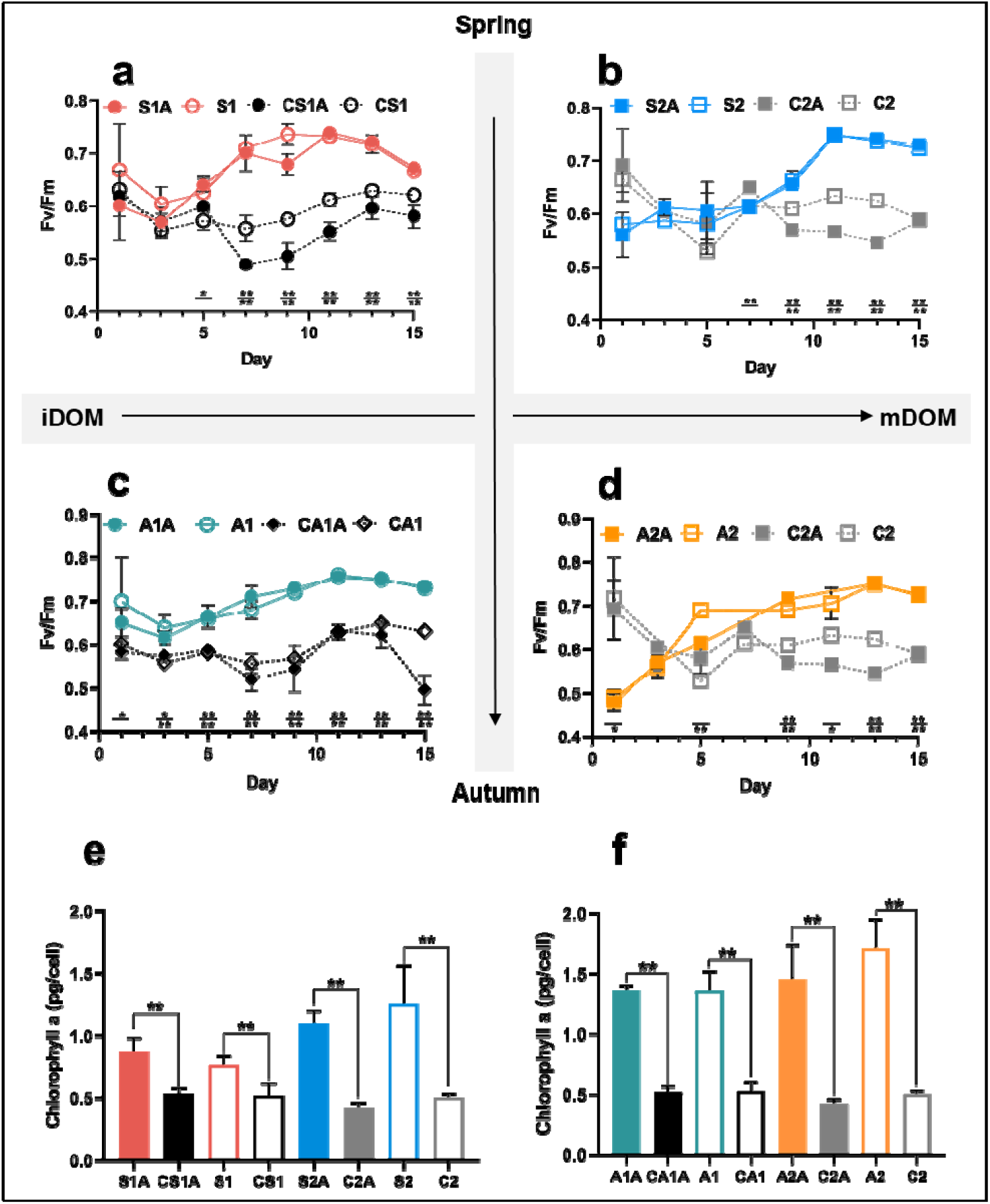
Fv/Fm and chlorophyll a content. (a) Fv/Fm under spring-iDOM conditions. (b) Fv/Fm under spring-mDOM conditions. (c) Fv/Fm under autumn-iDOM conditions. (d) Fv/Fm under autumn-mDOM conditions. (e)&(f) Average cellular chlorophyll a content of each group from 9th to 11th day. Data are the mean of biological triplicates with the error bar indicating standard deviation (mean ± SD, N=3).

### 3.2 Differences between iDOM and mDOM and seasonal variation

Pairwise comparison between iDOM and mDOM groups revealed that *in-situ* microbial degraded DOM (spring sampling) possessed a higher initial DIN and DIP (Supplementary Fig. 1a, c) and performed better in the promotion of cell growth with ∼47.3% (S2A vs. S1A) and 104% (S2 vs. S1) increase in the cell density (Fig. 4a). But in the autumn group, iDOM culture A1A performed better with ∼35.8% increase in the cell density than that in A2A (Fig. 4a), with significant higher initial DIN in A1A and A1(Supplementary Fig. 1d). The results of cellular Chla content were more consistent among the groups. All the mDOM groups showed an increase of 6.9% - 64.3% in cellular Chla content during the exponential growth phase compared to the iDOM groups, and with significant differences between S2 and S1, A2 and A1 (ANOVA, *P* <0.05) (Fig. 4b).

**Fig. 4.**
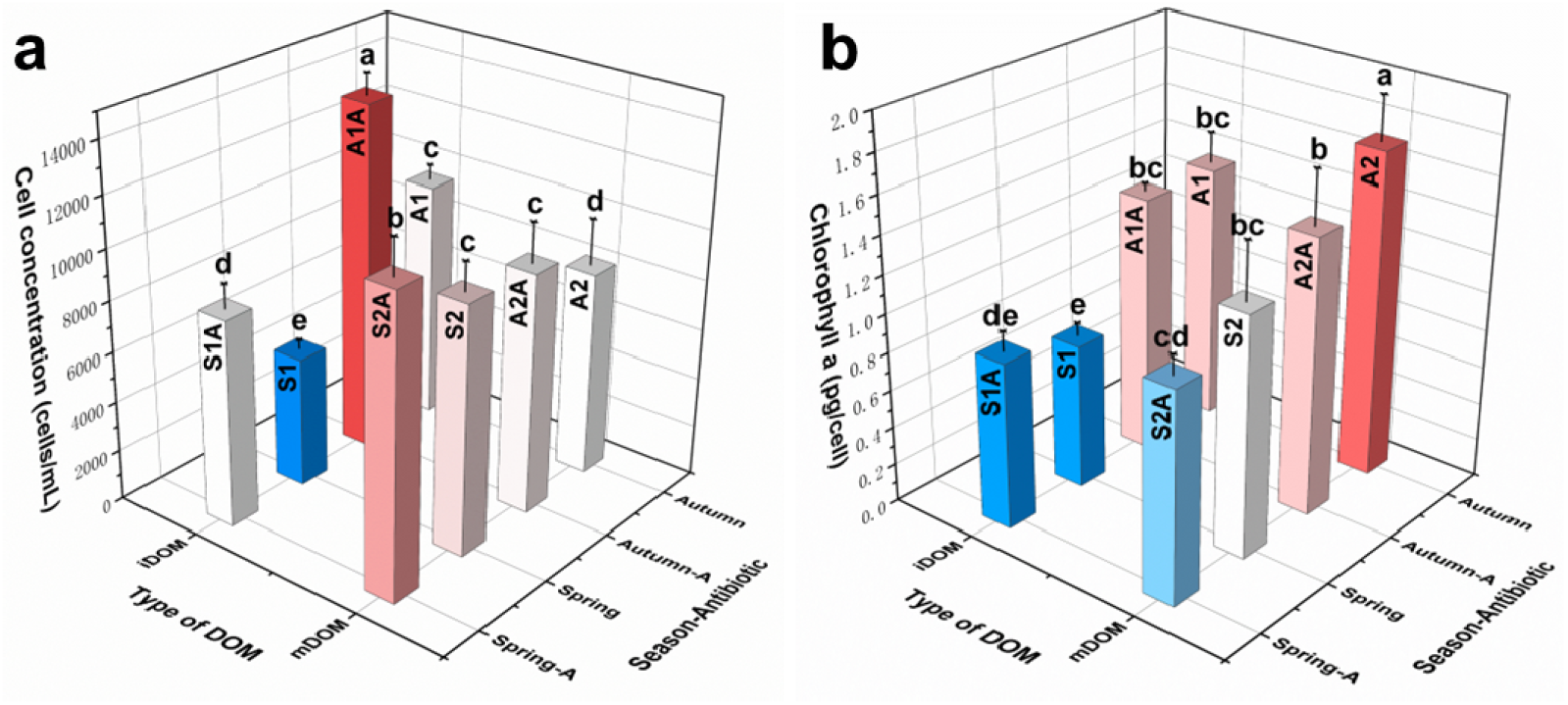
(a) Average cell concentration (Day 11∼15) under different DOM culture conditions (b) Average cellular chlorophyll a content (Day 9∼11) under different DOM culture conditions. Data are the mean of biological triplicates with the error bar indicating standard deviation (mean ± SD, N=3). Different letters above the columns indicate significant differences (ANOVA, P < 0.05).

Except for the variable effects of iDOM and mDOM, our data also showed seasonal differences in DOM. *P. donghaiense* under autumn-iDOM conditions had significantly higher cell densities (Fig. 4a) and notably increased growth rates (Fig. 2e). On the contrary, there was a stronger promotion of cell growth in spring-mDOM groups (Fig. 4a). In addition, the cellular Chla content was markedly elevated in all autumn-DOM groups. A1A and A1 improved by 55.6% and 77.7% over S1A and S1, respectively, while A2A and A2 only increased by 32.7% and 36.6% (Fig. 4b).

### 3.3 Cellular carbon, nitrogen, phosphorus

In the control group, average cellular C/N/P maintained stable throughout the experiment (Fig. 5a-c), and the median values of C/N and C/P were close to the empirical Redfield ratio (Fig. 5 d, e). Whereas in the DOM cultures, these values exhibited significant variation with regards to different DOM and the growth stages (Fig. 5). Compared with the control group, cells grown in the autumn DOM possessed higher cellular C (7.7%-36.6% increase), lower cellular N and P (53.7%, 26.3% decrease respectively), resulting in higher C/N and C/P values than Redfield ratio. In the contrary, lower cellular C (15.7%-34.8% decrease) with higher cellular N and P (84.9%, 48.3% increase respectively) were identified in cells grown in the spring DOM, as a result, both C/N and C/P values were slightly lower than the Redfield ratio but N/P ratio was significantly higher than the other groups and the Redfield ratio (Fig. 5d-f). Compared to autumn-DOM groups, in spring-DOM groups, there was a highly significant decrease in the cellular carbon content by ∼40%, but with a 2.68-fold higher nitrogen content and at least 35% greater phosphorus content (Fig. 5a-c).

**Fig. 5.**
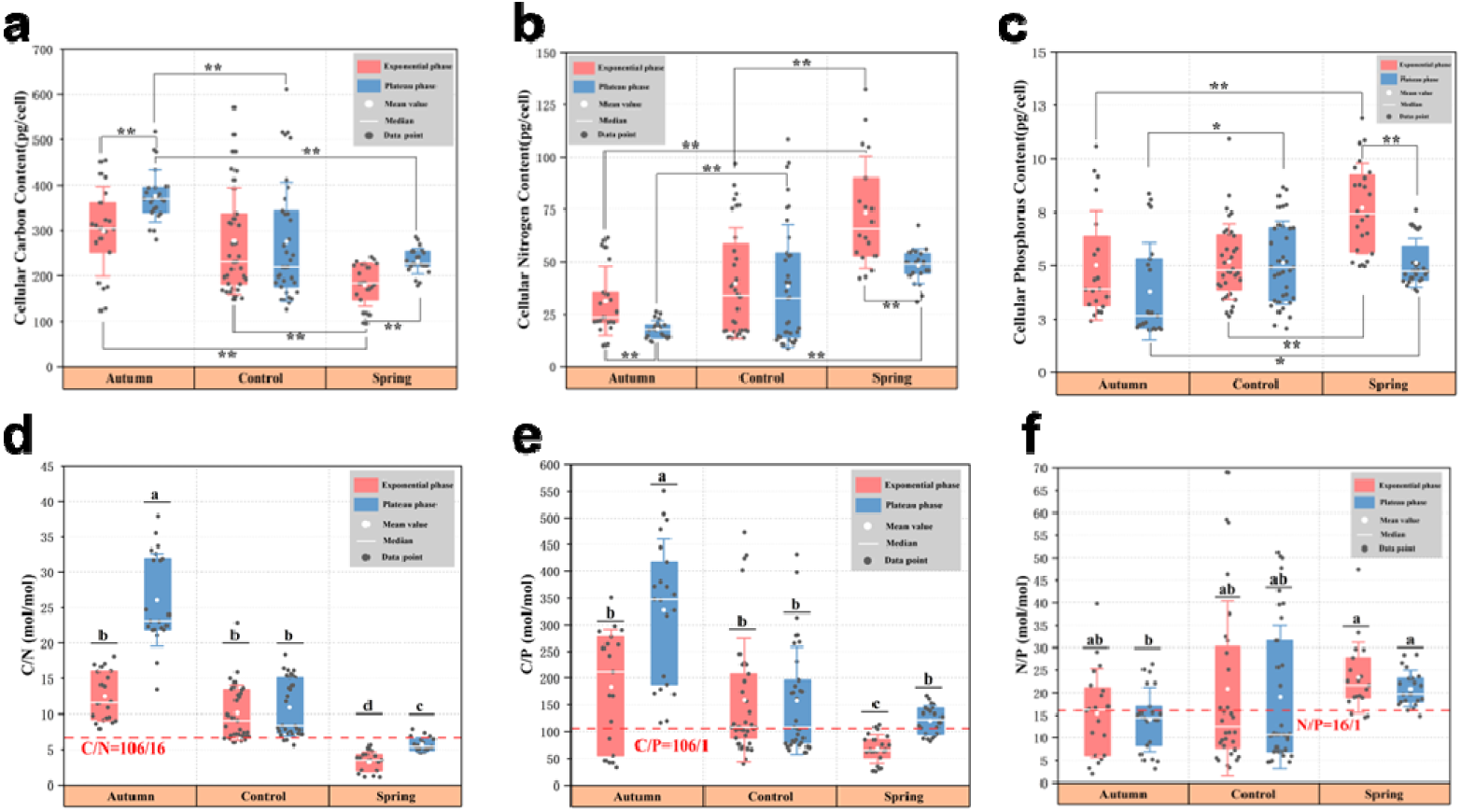
Cellular carbon, nitrogen, phosphorus content and their ratios during the exponential and plateau phases. (a) Cellular carbon content. (b) Cellular nitrogen content. (c) Cellular phosphorus content. (d) C/N ratio. (e) C/P ratio. (f) N/P ratio. P-value was calculated by T-test and ANOVA to represent significant difference between different groups. * P < 0.05, ** P < 0.01 and different letters above the columns also indicate significant differences (ANOVA, P < 0.05). Detailed data are provided in Supplementary Table. 2.

### 3.4 Comparison of cellular stoichiometry between DOM cultures and various culture conditions

To further examine the effect of *in-situ* DOM on the cellular stoichiometry, we collected published data of *P. donghaiense* grown under various culture conditions (Fig. 6). Upon comparison with other studies, we found that cells cultured with spring-DOM had a relative higher cellular carbon content than in other nitrogen-sufficient conditions (Fig. 6a), as well as significantly greater cellular nitrogen content with a low initial DIN (Fig. 6b). Overall, the C/N ratio in the spring scenario was lower than other studies, and the ratio in the autumn-DOM groups was notably higher than most studies (Fig. 6c). Given the fact that a considerable part of cell aggregates settles directly to the bottom after the bloom demise (Smetacek et al., 2012; Smith and Trimborn, 2024), we also compared cellular C/N ratio with reported value in surface sediments from different bloom areas, showing similar values between the spring-DOM culture and adjacent waters (Fig. 6d).

**Fig. 6.**
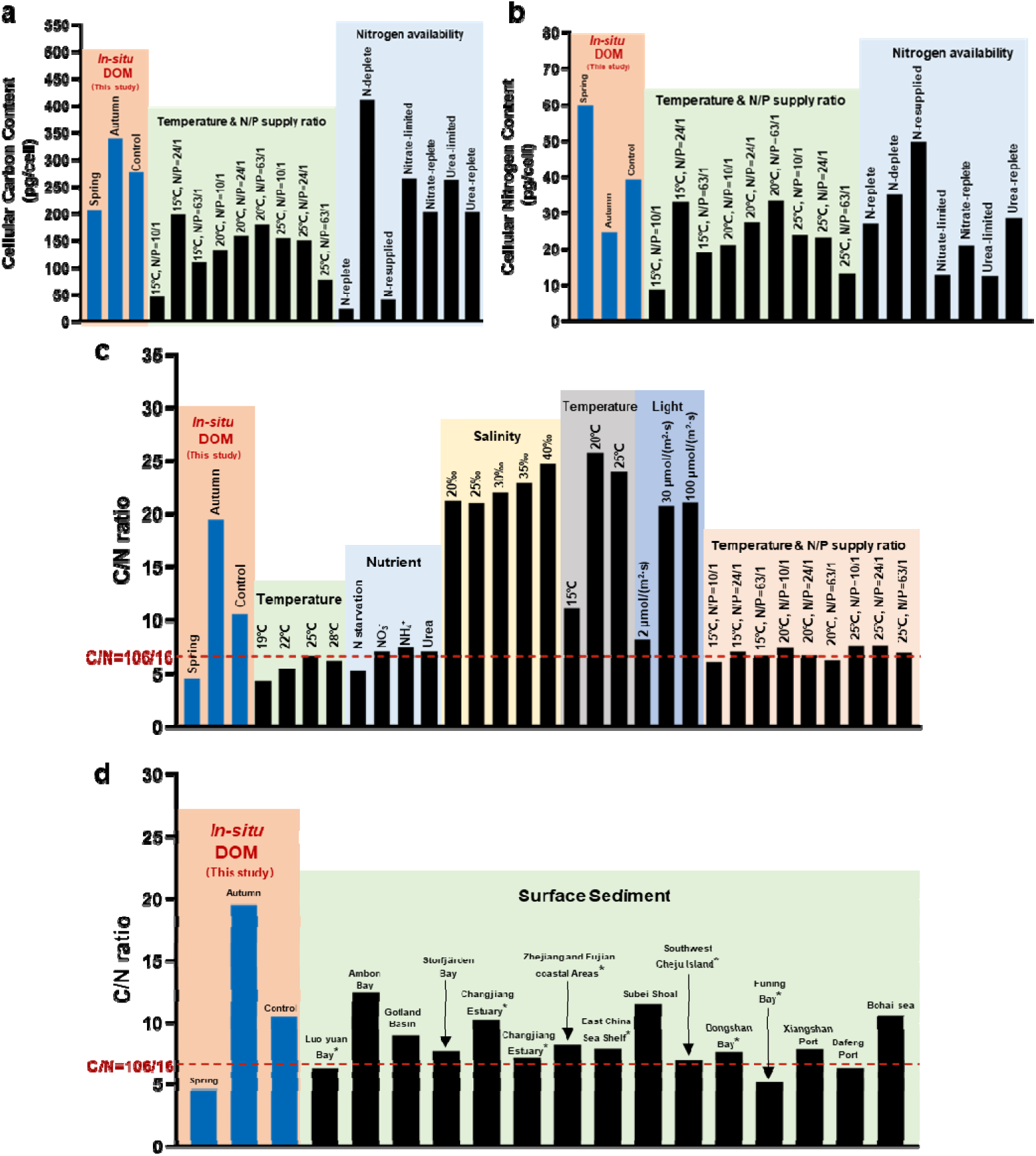
The comparison of cellular carbon and nitrogen content and the C/N ratios of P.donghaiense in this study with other studies. (a) Cellular carbon content (Zhang et al., 2015; Jing et al., 2017; Chen et al., 2019). (b) Cellular nitrogen content (Zhang et al., 2015; Jing et al., 2017; Chen et al., 2019). (c) C/N ratios (Cai et al., 2016; Liu, 2018; Chen et al., 2019; Zhang et al., 2022). (d) C/N ratios in this study and of surface sediments in some bloom areas (Hu et al., 2014; Chao et al., 2017; Wang et al., 2018; Rodil et al., 2020; Wang, 2020b; Likumahua et al., 2021; Xu, 2022; Beltran-Perez et al., 2023). The text above the column indicates different experimental conditions or areas. *: adjacent areas of this study (Fujian or East China sea). Detailed data are provided in Supplementary Table. 3 & 4.

## 4. Discussion

### 4.1 Promoted cell growth by mariculture DOM

The composition of DOM pool in mariculture waters is complicated and changing dynamically (Bouwman et al., 2013; Li, 2018), including small molecules (such as amino acid and urea) and not fully resolved organic matter enriched with N and P (Chen et al., 2023). The ability to utilize small organic molecules (ATP, urea etc.) has been extensively studied in *P. donghaiense* (Lin et al., 2016; Lu et al., 2022). The other organic matters can also be an alternative nutrient supply, especially for mixotrophic dinoflagellates which is able to digest the intact cell of the prey (Zhang et al., 2011; Jeong et al., 2021;). Our previous work revealed clathrin-mediated endocytosis in the uptake and utilization of DOP by *Phaeodactylum tricornutum* (Shu et al., 2022), and this mechanism was also recently reported in Symbiodiniacean species (Dinophyta) (Li et al., 2023). Besides organic nitrogen and phosphorus, dissolved metal ions (such as bioactive iron and copper), vitamins, and growth-promoting factors are accessible to dinoflagellates as well (Laglera and Berg, 2009; wells et al., 2015; Amin et al., 2015; Seymour et al., 2017). Thus, we can reasonably infer that mariculture DOM is able to sustain the outbreak of bloom by providing a comprehensive nutrient recipe for the rapid growth of mixotrophic dinoflagellates.

In this study, mariculture DOM served as an effective nutrient source promoting the growth of bloom forming species *P. donghaiense*, the growth rate was elevated 87.2% in comparison with the control group supplied with inorganic nutrients (Fig. 2). After 15 days culture, the cell density in the DOM culture is comparable with the average record of the bloom events (Zhang et al., 2019; Yu et al., 2020). It is consistent with relevant field investigation that dissolved organic nutrients were a crucial factor for the dominance of dinoflagellates (Zhang et al., 2023). Furthermore, for the first time, our results provide the quantitative evidence to address the question that *in-situ* DOM play an important role in initiating and sustaining the dinoflagellate blooms (Park et al., 2022). Our results also provide clues to the underlying mechanism to address the frequent *P. donghaiense* bloom in Changjiang Esturay, which is intensively affected by riverine discharge containing complex nutrient compositions in organic forms (Zhou et al., 2022; Wang et al., 2023).

Besides that, autumn-DOM exhibited a higher promotion of cell growth (Fig. 2c, d). Considering the projected warming scenario in the future, we propose a hypothesis that the outburst of HAB event in early autumn is likely to happen in ECS coastal waters, which may contribute to higher frequency of HABs shown in the model simulation (Xiao et al., 2018; Xiao et al., 2019; Zhou et al., 2022).

### 4.2 Different role of free-living and phycosphere microbiome in DOM transformation

To address the interaction between microbes and algae, the free-living community in the surrounding waters and phycosphere community are considered to function differently (Grossart, 2010; Stocker, 2012). Free-living microbiota are the main contributors to the degradation and re-mineralization of DOM, and decomposed small molecules are more accessible to microalgae (Kujawinski, 2011; Teeling et al., 2012; Chen et al., 2023). Our results are consistent with this scenario, both higher cell density and active photosynthesis activity (Fig. 2, Fig. 3) were observed in mDOM culture. Additionally, we noticed seasonal variation in the degradation of DOM by the *in-situ* microbiota, as evidenced by the different facilitation effects on cell growth, indicating the microbial community structure should be further investigated to elaborate the remineralization of DOM.

The interaction with the phycosphere microbiota and host is diverse, including mutualism, commensalism, antagonism, parasitism, and competition (Amin et al., 2012; Seymour et al., 2017). In our study, we observed that the antibiotic-treated groups exhibited slightly higher average growth rates in the exponential phase (Fig. 2e) and significantly higher cell densities (Fig. 2a, b, c, d), suggesting that there is a competition for nutrients in the presence of phycosphere bacteria. Considering that seed cultures were subjected to depletion of intracellular nutrient storage prior to the experiment, both algal cells and phycosphere bacteria were nutrient-stressed. In this case, both microalgae host and phycosphere bacteria could be competitive for resources (Thingstad et al., 1993, Joint et al., 2002). Opportunistic bacteria have been shown to be effective competitors for organic nitrogen compounds in coastal bloom of *Akashiwo sanguinea* (Han et al., 2021).

### 4.3 Cellular stoichiometry and implication on biogeochemical cycle

The initial DIP concentration of the experimental groups was similar, whereas the autumn-DOM groups had an obviously higher DIN concentration (Supplementary Fig. 1), yet the cellular nitrogen and phosphorus content markedly increased in the spring-DOM groups. Elevated N/P has been proposed to be a critical factor in the succession of diatoms to dinoflagellates during the bloom (Li et al., 2021b; Lu et al., 2022). Our autumn-iDOM cultures are in line with this perspective, yet most *in-situ* DOM groups with low initial N/P ratio (Supplementary Fig. 1e) was also able to stimulate the cell growth significantly. These results suggest that dissolved organic matter play a key role in the outburst and maintenance of the HAB events. We should also point out that, in this experiment, we only quantify the promotion in batch culture; however, in practice, it should be taken as a continuous culture system, where a higher promotion effect is expected. We propose that model simulation should be introduced to such a scenario for further elaboration.

Compared to the biological pump in the open ocean, most HABs occur in shallower coastal waters, and massive cells may form high-density particulate matter and settle directly to the bottom without adequate time for microorganisms to decompose, leading to frequent bottom hypoxia and altering the elemental flux ratios to depth (Li et al., 2019; Burdick et al., 2020; Smith and Trimborn, 2024). Thus, it’s essential to investigate the cellular elemental stoichiometric ratios, which are involved with the uptake and transformation of mariculture DOM and pose a cascade effect on the biogeochemical cycle in adjacent waters and sediments (Fig. 7). Comparison of cellular stoichiometry implies that during spring HAB events in coastal mariculture areas, more organic carbon, nitrogen, and phosphorus may be transported to the bottom layer along with the cell settling. And relatively low C/N and high N/P ratios of algal cells might pose effect on the elemental composition and stoichiometry of surface sediments.

**Fig. 7.**
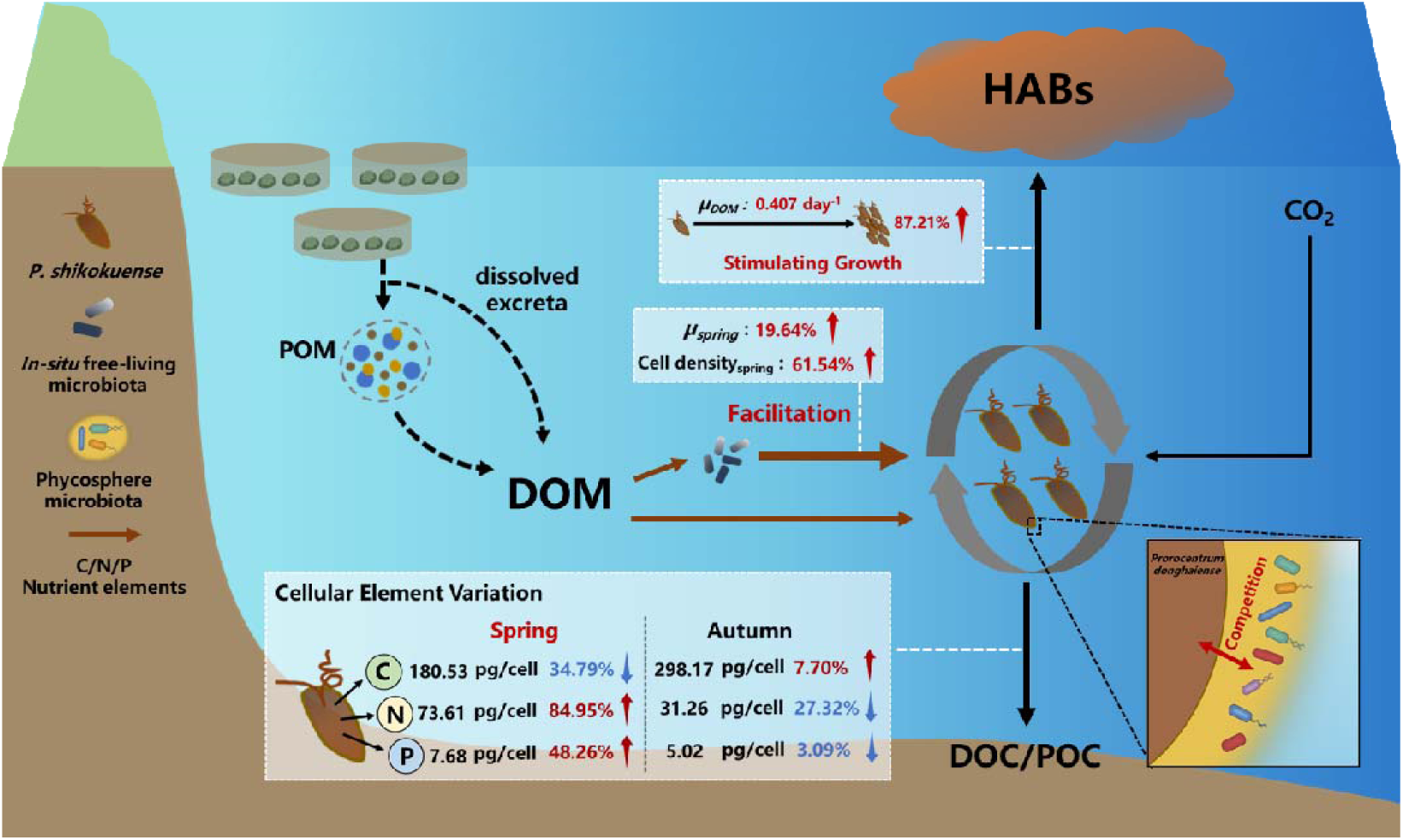
Schematic diagram of the promoted cell growth by in-situ DOM and ecological implications in biogeochemical cycle. Red numbers and arrows indicate increases, while blue ones represent decreases.

Compared to control groups, our results showed a significant variation in cellular carbon, nitrogen, and phosphorus content that differed between the spring and autumn DOM groups, exhibiting different nutrient uptake strategies (Fig. 5, 7). In the spring-DOM groups, we observed a significant elevation of cellular N and P content compared with control groups, suggesting a higher supply of organic N and P compounds in the surrounding waters, and the accumulation of cellular carbon was constrained, likely due to the extra energy cost in the assimilation of organic nutrients (Falkowski, P. G. and Raven, J. A., 2007). In comparison, in the autumn-DOM groups, we observed significantly higher cell density, cellular C, along with decreased cellular N and P content, suggesting a potential contribution of DOC to the rapid growth. Taking these results together, metabolites from mariculture species during the life cycle are suggested to be introduced into water quality monitoring.

## Conclusion

We use dissolved organic matter collected from mariculture area as the sole medium to investigate the DOM utilization capacity of *P. donghaiense*, a typical HAB causative species in ECS. Our results quantified the efficient utilization of *in-situ* DOM (markedly increased cell density, average growth rate, photosynthetic efficiency, and cellular Chla content) and its significant effect on the cellular element content and stoichiometry. Meanwhile, different sets of parallel treatments revealed the facilitation conferred by *in*-*situ* microbiota in DOM remineralization, while there may be competitive or exploitative interactions between phycosphere microbiota and dinoflagellates when resources are scarce. Our results provided rigorous evidence that dissolved organic matter might play a more vital role in the costal bloom events than previously suggested. The ability to efficiently utilize dissolved organic matter is a key strategy adopted by mixotrophic *P*. *donghaiense* to outcompete other phytoplankton species in organic matter-rich waters. The assimilation of DOM greatly shapes the cellular stoichiometry and might have a profound effect on the biogeochemical cycle of key elements in mariculture ecosystems.

## CRediT authorship contribution statement

**Hongwei Wang**: Methodology, Investigation, Formal analysis, Writing – original draft, Writing - Review & Editing. **Jian Ma**: Methodology, Writing - Review & Editing. **Siyang Wu**: Investigation. **Yiting Hong**: Investigation. **Chentao Guo**: Methodology**. Jing Zhao**: Writing - Review & Editing. **Xin Lin**: Conceptualization, Methodology, Writing - Review & Editing, Funding acquisition, Supervision.

## Data availability

Data will be made available on request.

## Acknowledgements

This study was supported by the National Key R&D Program of China (2022YFC3105302) and State Key Laboratory of Marine Environmental Science Internal Program No. MELRI2302.

We thank Weiwei You and the lab members (Xiamen University) for helping us collect the mariculture seawater, Junhui Chen (Xiamen University) for helping with carbon and nitrogen determination and Ling Li, Yujie Wang and Marine Ecogenomic Lab (MEG) for the technical assistance in the experiment. We are also grateful to Tengyue Fang (Xiamen University) for helping nutrient determination.

