## Supplementary Table.1 for "Stimulated *Prorocentrum donghaiense* cell growth by *in-situ* mariculture dissolved organic matter"

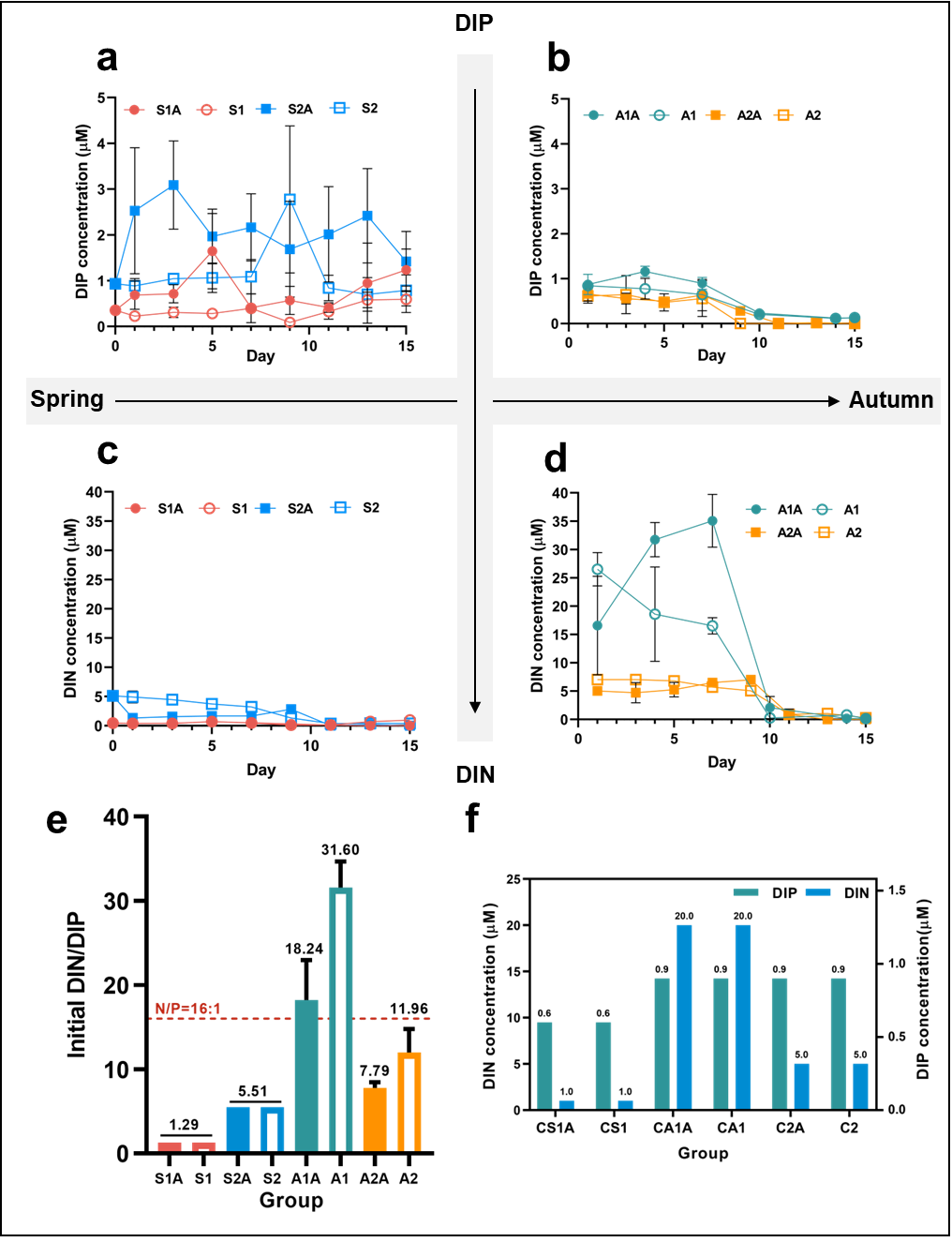

Supplementary Fig.1 Changes in inorganic nutrients during the culture. (a) DIP under spring-DOM conditions. (b) DIP under autumn-DOM conditions. (c) DIN under spring-DOM conditions. (d) DIN under autumn-DOM conditions. (e) Initial DIN/DIP in each group. (f) Initial DIN and DIP in control groups.

Supplementary Table.1 Experimental settings of culture groups (* means the DIP and DIN concentration was measured on day 1 instead of day 0)

| **Group** | **Season** | | **Filtration** | | **Antibiotic** | | ***in-situ* Nutrient (µM)** | |
| --- | --- | --- | --- | --- | --- | --- | --- | --- |
|  | **Spring** | **Autumn** | **iDOM (1)** | **mDOM (2)** |  |  | **DIP** | **DIN** |
| **S1A** | **+** |  | **+** |  | **+** |  | **0.35** | **0.46** |
| **S1** | **+** |  | **+** |  |  | **+** | **0.35** | **0.46** |
| **S2A** | **+** |  |  | **+** | **+** |  | **0.93** | **5.14** |
| **S2** | **+** |  |  | **+** |  | **+** | **0.93** | **5.14** |
| **A1A** |  | **+** | **+** |  | **+** |  | **0.871*** | **16.58*** |
| **A1** |  | **+** | **+** |  |  | **+** | **0.839*** | **26.52*** |
| **A2A** |  | **+** |  | **+** | **+** |  | **0.656*** | **5.04*** |
| **A2** |  | **+** |  | **+** |  | **+** | **0.613*** | **7.04*** |
| **CS1A** | **Artificial seawater**  **(Control group)** | | | | **+** |  | **0.6** | **1** |
| **CS1** |  |  |  |  |  | **+** |  |  |
| **CA1A** |  |  |  |  | **+** |  | **0.9** | **20** |
| **CA1** |  |  |  |  |  | **+** |  |  |
| **C2A** |  |  |  |  | **+** |  | **0.9** | **5** |
| **C2** |  |  |  |  |  | **+** |  |  |

Supplementary Table.2 Cellular carbon, nitrogen, phosphorus content and C/N, N/P and C/P ratio

（exp: exponential phase; pla: plateau phase）

| **Group** | | **Carbon**  **(pg/cell)** | **Nitrogen (pg/cell)** | **Phosphorus (pg/cell)** | **C/N** | **C/P** | **N/P** |
| --- | --- | --- | --- | --- | --- | --- | --- |
| **Spring** | **exp** | **180.53±47.82** | **73.61±26.84** | **7.68±2.10** | **3.25±1.45** | **67.46±26.69** | **23.35±7.82** |
|  | **pla** | **232.70±28.41** | **48.04±8.27** | **5.13±1.13** | **5.68±0.96** | **120.95±27.25** | **20.93±4.04** |
| **Autumn** | **exp** | **298.17±99.16** | **31.26±16.48** | **5.02±2.55** | **12.53±3.47** | **184.15±107.03** | **15.49±9.84** |
|  | **pla** | **375.15±58.44** | **17.87±4.28** | **3.79±2.28** | **26.10±6.55** | **326.93±133.59** | **14.13±7.13** |
| **Control** | **exp** | **276.86±117.92** | **39.80±26.54** | **5.18±1.78** | **10.17±3.97** | **159.04±115.27** | **20.95±19.49** |
|  | **pla** | **275.36±131.08** | **38.59±29.23** | **5.14±1.95** | **10.89±4.04** | **158.13±100.28** | **19.09±15.96** |

Supplementary Table.3 Cellular carbon, nitrogen content and C/N ratio of *P. donghaiense* in this study and other studies. (-: No data)

| **Treatment** | | **Cellular Carbon Content (pg/cell)** | | **Cellular Nitrogen Content (pg/cell)** | **C/N Ratio** | **Ref** |
| --- | --- | --- | --- | --- | --- | --- |
| **DOM** | **Spring** | **206.62** | | **59.84** | **4.53** | **This study** |
|  | **Autumn** | **338.93** | | **24.41** | **19.48** |  |
|  | **Control** | **276.11** | | **39.17** | **10.53** |  |
| **Temperature** | **19℃** | **-** | | **-** | **4.34** | **Zhang et al., 2022** |
|  | **22℃** | **-** | | **-** | **5.48** |  |
|  | **25℃** | **-** | | **-** | **6.68** |  |
|  | **28℃** | **-** | | **-** | **6.24** |  |
| **Nutrient** | **N starvation** | **-** | | **-** | **5.25** | **Liu, 2018** |
|  | **NO_3_-** | **-** | | **-** | **7** |  |
|  | **NH_4_+** | **-** | | **-** | **7.39** |  |
|  | **Urea** | **-** | | **-** | **7** |  |
| **Salinity** | **20‰** | **-** | | **-** | **21.26** | **Cai et al., 2016** |
|  | **25‰** | **-** | | **-** | **20.95** |  |
|  | **30‰** | **-** | | **-** | **22.01** |  |
|  | **35‰** | **-** | | **-** | **22.87** |  |
|  | **40‰** | **-** | | **-** | **24.67** |  |
| **Temperature** | **15℃** | **-** | | **-** | **11.06** |  |
|  | **20℃** | **-** | | **-** | **25.75** |  |
|  | **25℃** | **-** | | **-** | **23.97** |  |
| **Light** | **2 μmol/(m^2^*s)** | **-** | | **-** | **8.15** |  |
|  | **30 μmol/(m^2^*s)** | **-** | | **-** | **20.71** |  |
|  | **100 μmol/(m^2^*s)** | **-** | | **-** | **21.01** |  |
| **15℃** | **N:P supply ratio=10:1** | | **190.3** | **32.7** | **6.13** | **Chen et al., 2019** |
|  | **N:P supply ratio=24:1** | | **197.3** | **33.7** | **6.99** |  |
|  | **N:P supply ratio=63:1** | | **150.3** | **28** | **6.74** |  |
| **20℃** | **N:P supply ratio=10:1** | | **209** | **32.7** | **7.35** |  |
|  | **N:P supply ratio=24:1** | | **151** | **24** | **6.75** |  |
|  | **N:P supply ratio=63:1** | | **156.7** | **23.3** | **6.30** |  |
| **25℃** | **N:P supply ratio=10:1** | | **128** | **20.3** | **7.58** |  |
|  | **N:P supply ratio=24:1** | | **140.5** | **23.5** | **7.59** |  |
|  | **N:P supply ratio=63:1** | | **138** | **21** | **6.88** |  |
| **Nitrogen availability** | **N-replete** | | **23.71** | **27** | **-** | **Zhang et al., 2015** |
|  | **N-deplete** | | **410.687** | **35** | **-** |  |
|  | **N-resupplied** | | **40.41** | **49.67** | **-** |  |
|  | **Nitrate-limited** | | **264.48** | **12.64** | **-** | **Jing et al., 2017** |
|  | **Nitrate-replete** | | **202.13** | **20.905** | **-** |  |
|  | **Urea-limited** | | **262.3** | **12.33** | **-** |  |
|  | **Urea-replete** | | **201.905** | **28.435** | **-** |  |

Supplementary Table.4 C/N ratios of surface sediments in some bloom areas. (*: adjacent areas of this study, Fujian or East China sea)

| **Location** | **C/N ratio (Average)** | **Ref** |
| --- | --- | --- |
| **Luoyuan Bay*** | **6.21** | **Wang et al., 2018** |
| **Ambon Bay** | **12.38** | **Likumahua et al., 2021** |
| **Gotland Basin** | **9** | **Beltran-Perez et al., 2023** |
| **Storfjärden Bay** | **7.72** | **Rodil et al., 2020** |
| **Changjiang Estuary*** | **10.24** | **Chao et al., 2017** |
| **Changjiang Estuary*** | **7.2** | **Hu et al., 2014** |
| **Zhejiang and Fujian coastal areas*** | **8.2** | **Xu, 2022** |
| **East China Sea Shelf*** | **7.9** |  |
| **Subei Shoal** | **11.5** |  |
| **southwest Cheju Island*** | **7** |  |
| **Dongshan Bay*** | **7.64** | **Wang, 2020** |
| **Funing Bay*** | **5.18** |  |
| **Xiangshan Port** | **7.86** |  |
| **Dafeng Port** | **6.22** |  |
| **Bohai sea** | **10.6** |  |
